# Seed Germination, Post-fire Plant Growth and Conservation of the Rare Endemic and Endangered *Chamaecrista glandulosa* var. *mirabilis* (Fabaceae)

**DOI:** 10.1101/2022.03.04.483036

**Authors:** Fernando J. Vilá Terrada, Jonathan A. López Colón

## Abstract

Conservation has been challenged by biodiversity loss drivers. Also, fire disturbance can temporarily change ecosystems. Fire effects in soil nutrients and pH, plant abundance, reproduction, seed weight, seed germination, and plant growth were assessed. Four seed starting systems were used for germination and growth under laboratory conditions. We found significant differences in the means of iron, manganese, nickel, soil pH, and plant height, and a significant positive linear relationship between seed weight and plant height. Results indicate that fires increase soil pH and cause changes in micronutrients that can increase plant growth. Large size plants are produced from high weight seeds. Finally, *ex situ* conservation and species reintroduction were feasible conservation strategies that should be integrated with *in situ* conservation.

## Introduction

Ecosystems are sensitive to natural and anthropogenic disturbances (Adeney et al. 2016) that change biodiversity. Endangered rare endemic species are of priority for conservation due to its restricted and narrow geographic distribution, small abundance, and critical state. Conservation has been challenged by the increasing biodiversity loss drivers like habitat loss and change, overexploitation, pollution, invasive species, and climate change (Heywood 2017). Conservation biology addresses the biology of species, communities, and ecosystems that are perturbed with the goal of providing principles for preserving and protecting biodiversity (Soulé 1985). Plant conservation is dependent on the creation of protected areas and parks and is complemented by conservation strategies and research in population biology (Soulé 1985, Heywood 2017), including environmental conditions experiments.

Fire disturbance has the ability to temporarily produce significant changes to organic matter, chemical properties, and ecosystems (Bridges et al. 2019). Natural charcoal and ash from wildfire have the potential to alter the soil solution chemistry (Gundale and DeLuca 2006). Plant available nitrogen and total phosphorus concentrations were found to increase in burned soils (Kutiel and Naveh 1987, Paritsis et al. 2006), but decreases in exchangeable potassium (Bridges et al. 2019), dissolved nitrogen and soil moisture (Knelman et al. 2017) have also been found. Specifically, increases in available calcium, magnesium, and phosphorus have resulted from high intensity fires (Bridges et al. 2019), and decreases in carbon and nitrogen concentrations were caused by high fire frequencies (Pellegrini et al. 2018). Advances in flowering and fruiting onset and greater fruit set (Paritsis et al. 2006), along increases in shoot and root biomass, seed number, and seed weight (Kutiel and Naveh 1987) have resulted in burned environments.

Growth, irreversible increase in size, form, or number with time (Hunt 2003), can have implications for the life cycle of plants. The growth and distribution of plants are controlled by the interaction of gene mechanisms and the environment, which includes factors like radiation, temperature, water, atmospheric gases, soil chemistry, and biotic factors, among others (Billings 1952). Plant size was found to be positively correlated with absolute growth (McIntosh 2002), positively correlated with the number of buds, flowers, and fruits produced by plant (Rojas-Sandoval and Meléndez-Ackerman 2011), and positively related to inflorescence size (Rodríguez-Robles et al. 1992). Also, greater plant size (Dollard 2018) and absolute plant and root mass (Schreeg et al. 2005) may be related with high survival. Therefore, greater plant growth and size promotes reproduction and survival that can influence population size and the preservation of species.

Seeds are an important source for plant propagation. Seed weight was found to be positively correlated with plant size (Hendrix 1984), and similarly, seed mass was positively correlated with shoot mass, root mass, and shoot length (Mukherjee et al. 2019). Large seed weight showed higher germination percentage (Anjusha et al. 2015), and high seed weight and width resulted in faster germination (Xu et al. 2016). Seed germination is the process of seedling formation that initiates with the uptake of water (imbibition), the reactivation of cellular and metabolic processes, and the extension of the radicle or embryonic root (Bewley 1997). Abiotic factors that influence seed germination include soil pH (Singh et al. 1975), light, temperature, soil type, and sowing depth (Shen et al. 2015).

Main conservation strategies for plants are *in situ* and *ex situ* conservation and species reintroduction (Ren et al. 2014). Propagation from seeds was found a viable method for *ex situ* conservation (Cerabolini et al. 2004). Species reintroduction is the bridge between *in situ* and *ex situ* conservation and is the final goal of *ex situ* conservation (Ren et al. 2014). Augmentation, a main concept of reintroduction, refers to population reinforcement when individuals are added to an existing population increasing the population size or genetic diversity (Ren et al. 2014). The use of seedlings and mixing material from diverse populations promoted higher survival rates in reintroductions (Godefroid et al. 2011). Finally, transplants as founders yielded greater establishment rates than seeds (Guerrant Jr and Kaye 2007).

Studies on *ex situ* plant development and field establishment following transplantation are required for species reintroductions (Cerabolini et al. 2004). Field observations suggested differences in population size and plant growth between populations of the study species. The main objectives were to propagate from seeds, germinate in field and laboratory conditions, and test *ex situ* conservation and reintroduction as reinforcement. Fire effects in soil nutrients and pH, plant abundance, and reproduction in the field were examined. Also, fire effects in seed weight, seed germination, and plant growth in the laboratory were assessed. The first question was, what are the effects of fire on soil nutrients and pH? Second, what are the effects of post-fire soil nutrients and pH on the species? We hypothesized that soil nutrients and pH will increase after fire and positively affect seed weight, plant growth, reproduction, and plant abundance in the fire affected population. At last, is seed germination related to soil pH, and is plant height related to seed weight? We hypothesized that seed germination will be related to soil pH and plant height will be positively related to seed weight.

## Materials and Methods

### Study species

*Chamaecrista glandulosa* var. *mirabilis* (Pollard 1902) (Fig. 1a) is a rare endemic and endangered plant species in Puerto Rico. It belongs to the Fabaceae family and Caesalpinioideae subfamily. It is endemic to white silica sands of the north coast of Puerto Rico at elevations near sea level (USFWS 1994). The species was classified as endangered on April 5, 1990 according to the Endangered Species Act of 1973 (USFWS 2015). It is endangered due to sand extraction, deforestation, and urban and industrial expansion (USFWS 1994). Main threats include habitat destruction, habitat modification, climate change, invasive plant species, and human-induced fires (USFWS 2015).

**Figure 1.**
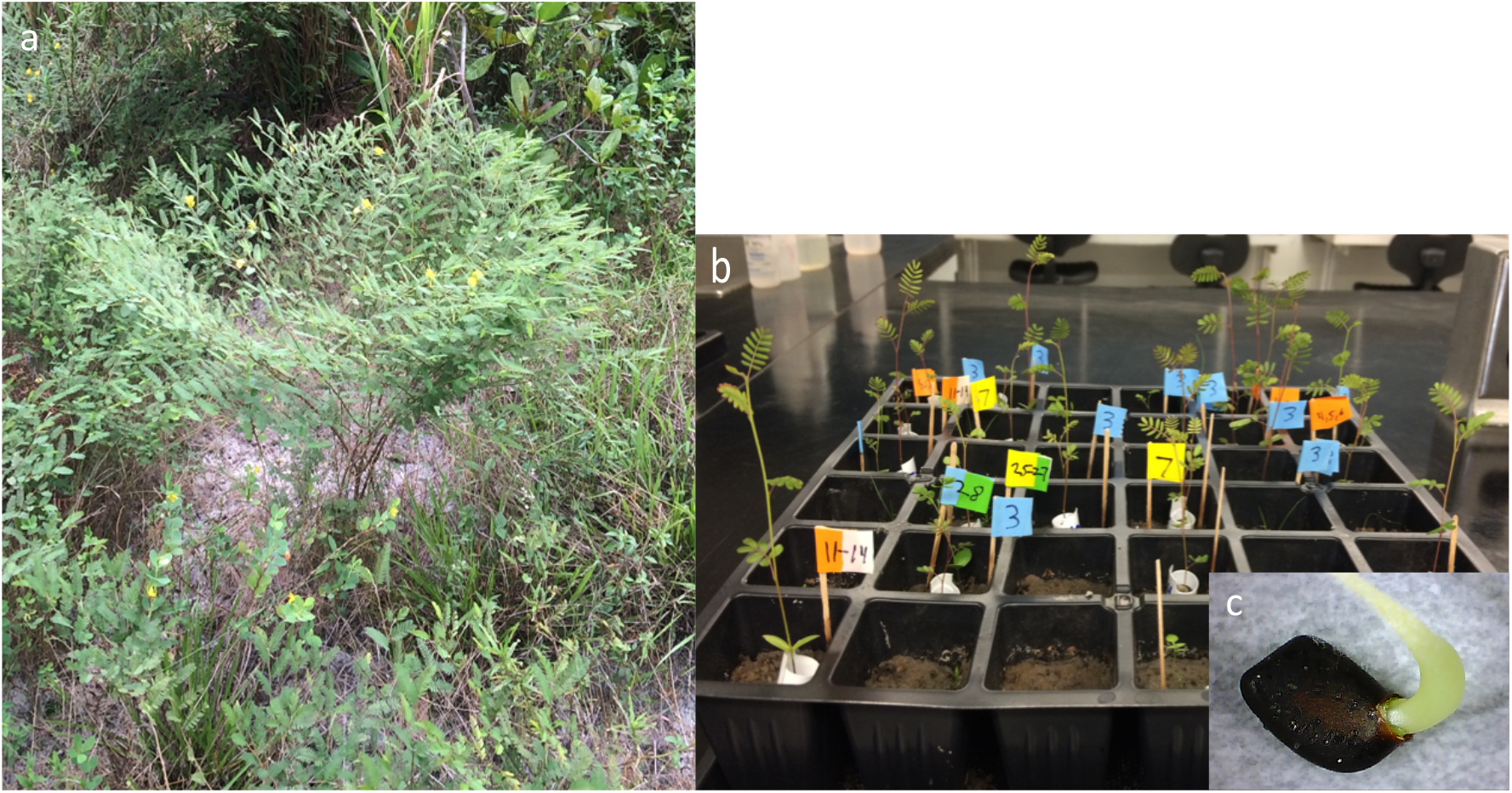
*Chamaecrista glandulosa* var. *mirabilis* (a). Plant growth in the systems (b). Seed germination in the laboratory (c).

The species is a small perennial shrub. The plant grows from 1-8 cm tall prostrated or assurgent. Leaves are compound paripinnate with alternate arrangement, measuring up to 4.5 cm long. Leaflets are linear-elliptic with 10-18 pairs, and measure up to 7.5 mm long and 1.4 mm broad. Solitary flowers have 3 short petals and a big long abaxial petal. The corolla is yellow. The fruit is a flat dehiscent legume, 18-40 mm long, 3-3.8 mm broad, and seeds are 2.2-2.9 mm long (Liogier 1988).

### Study site

Two populations were studied in a tropical coastal wetland at Reserva Natural Laguna Tortuguero (RNLT), located between the municipalities of Vega Baja and Manatí (18°27’48’’N, 66°26’29’’W) on the north coast of Puerto Rico. The wetland includes a marsh, swamp, two lagoons, and a dry evergreen or littoral forest (USFWS 1994) within the subtropical moist forest life zone (Miller and Lugo 2009). The soil is white silica (SiO_2_) sand with high acidity, extreme nutrient scarcity, and low organic matter content (Adeney et al. 2016). Puerto Rico has a warm and humid tropical climate. The mean annual air temperature ranges from 24°C to 27°C in coastal areas, and rainfall is about 1,825 mm in an average year (Gómez-Gómez et al. 2014).

Populations are located on open areas in the forest near the Tortuguero lagoon. Population 1 is located southeast of the lagoon in an open area. Population 2 is located south of the lagoon in a semi-open area with canopy and trails. A human-induced fire occurred in the area of Population 1 in May 2015, affecting the population and serving as an experiment. Experimental observations and tests were conducted in the laboratory and the field.

### Sampling and experimental design

Two hectares (Plot 1 and Plot 2) measuring 100 m x 100 m (10,000 m^2^) were established to sample the populations. The experimental design included two groups, the experimental and control. The experimental group was Population 1, which was affected by the fire. The control group was Population 2, which was not affected by fire. Nine replicates of 36 seeds by population were done for the seed germination rate and plant growth. Three were done for germination percent in sand and one for germination percent in different soils. Two replicates were used to sample seed weight and reproduction. And three replicates were used for field germination and transplantation.

### Soil samples

Soil samples were taken in both plots for soil chemistry analysis. Samples were collected in 3 different points as replicates within each plot and divided in 3 for a total of 18 samples (9 in Plot 1; 9 in Plot 2). All samples were collected using a 15 in long stainless-steel tubular sampler, which included the O (organic) and A (surface) horizons. After collection, the sand soil was divided by organic matter and surface layers and stored in plastic bags. The soil samples were taken to Environmental Quality Laboratories, Inc. for chemical tests after 2 years of the fire event.

### Fruit collection

Mature fruits were collected from adult plants in the field to assess reproduction differences between populations. A total of 107 fruits were collected in each population. They were stored in plastic bags and taken to the laboratory. There, fruits were opened, and seeds were counted per fruit by population. Seeds were stored in centrifuge tubes at ambient temperature. Seeds (n = 216) were identified with a number and weighed on a scale by population in the laboratory. Finally, seeds were tracked after sowing to determine seed germination and plant growth. At last, plant abundance was counted according to the development stage of plants (seedling, juvenile, adult) within both plots.

### Seed germination

Four Burpee – 36 Cell Seed Starting Systems were used for seed germination and plant growth under laboratory conditions as part of the experiment. Each system included: a 36-cell planting tray, self-watering mat, plant stand, watering tray, and a clear greenhouse dome. Of the four seed starting systems, two were used for the sample of Population 1 and the other two for the sample of Population 2. Silica sand of Plot 1 was added to the cells of planting trays of Population 1, and silica sand of Plot 2 was added to the cells of planting trays of Population 2. Seeds of Population 1 were sowed in the soil on the planting trays of Population 1, and seeds of Population 2 were sowed in the soil on the planting trays of Population 2. Then, a small amount of water was added in each cell with a dropper for imbibition. Finally, the clear greenhouse domes were placed on top of the planting trays for the first two weeks, and water was added to the watering trays.

The four seed starting systems were placed on bigger trays in a rack under a lamp containing two light bulbs (130v 150w 1,150LM). Yellow light from the light bulbs was provided during the day. Light bulbs were powered on every morning and powered off in the afternoon for the night. The photoperiod was established with 10 hours of light and 14 hours of darkness. Water was added to the watering trays every time they were empty to provide humidity. Seeds were monitored almost daily in each system since sowing day for 60 days. They were identified and marked in the cells with little flags made with toothpicks and colored tapes when germination occurred each day. Finally, germinated seeds were counted per day by population to perform the seed germination rate and percent.

Other two germination tests were conducted, seed germination in different soils and in the field. Seeds from both populations were sowed in pots with Pro-Mix potting and seeding mix with Peat moss, potting mix and sand in the surface, garden soil, and Burpee pellets to examine germination on different soils in the laboratory. Later, germinated seeds were counted. For seed germination in the field, three quadrants measuring 1 m x 1 m were made near Plot 1. Each quadrant was weeded first and then marked in the corners with flags. Sixteen seeds were sowed in the surface of the soil in each quadrant in the month of November. A total of 48 seeds were used, including seeds of both populations. Also, a plot measuring 3 m long x 2 m wide was established using flags near Plot 2. Forty-eight seeds of the species were sowed in the surface of the soil without vegetation in the month of March. At last, another plot measuring 5 m long x 2 m wide was made in a new open area to create a population from seeds. Seeds were sowed in soil without vegetation in the month of March. Water was added in the soil surface once in each plot for germination. Finally, seed germination was monitored for 5 months.

### Plant growth

An analysis using size was made to understand plant growth in the seed starting systems under laboratory conditions. The analysis was made by the classical quantitative method, which involves measurements through time (relatively infrequent long intervals) in a large number of plants (Hunt 2003). After 2 weeks since sowing day (day 15), seedling height was measured in cm by population with a metric ruler in each seed starting system. Every 2 weeks, juvenile plants were measured again. This was done for a total of 4 measurements until day 57. After the first measurement, each seedling was identified with a seedling tag and a number.

### Acclimation and reintroductions

Juvenile plants from the seed starting systems were transplanted to pots. Pots were placed in a nursery for acclimation within a metallic cage to protect them from herbivory of green iguanas. Plants were in the acclimation process for 2 weeks, and they were watered twice a week. In the nursery, air temperature was measured from 29-37°C and relative humidity from 46-60% using a hygrometer. Soil temperature was measured from 27-32°C with a soil thermometer.

After acclimation, plants were transplanted into the wild populations areas to perform species reintroductions as reinforcement. Two reintroductions were made for Population 1 and one for Population 2 for a total of three. The first transplant was made near Plot 1 in an area without vegetation inside 2 quadrants that measured 1 m × 1 m. Each plant was placed in the corners of the quadrants, except one that was placed in the middle. Both quadrants contained 5 plants for a total of 10 plants planted. The second transplant near Plot 1 was made inside a plot that measured 12 m long × 2 m wide. The area was weeded first and then marked with flags. A total of 16 plants were planted. The third transplant was made in an open area without vegetation in a plot that measured 3 m long × 2 m wide near Plot 2. This plot was marked with flags in the corners, and 3 plants were planted. Overall, 29 plants were reintroduced to the natural habitat. All individuals were identified with a metallic tag and a number, and plant survival was recorded monthly. Finally, the second transplant near Plot 1 was observed to examine the reproduction of reintroduced plants (n=16). Buds, flowers, and fruits were identified by plant weekly to determine the proportion of plants reproducing out of all plants alive.

### Data analysis

The statistical analysis of data was made in RStudio and XLSTAT using hypothesis tests and linear models. The two independent samples T test was used to compare means of soil nutrients and pH, plant abundance, and plant height between Population 1 and Population 2. The Mann Whitney U test was done to compare the distribution of soil nutrients and pH, seed germination, seed weight, and plant reproduction between Population 1 and Population 2. Finally, we tested linear regressions between seed weight and plant height and soil pH and seed germination percent.

## Results

### Soil chemistry

Soil nitrate and nitrite, nitrogen, phosphorus, and organic matter content between plots were found to be equal in means (*P*>0.05; *t*=-0.49 to 1.14; *df*=16). Copper and zinc concentrations among plots were found to be equal in distribution (*P*>0.05; *U*_cu_=28; *U*_zn_=32). We found statistically significant differences in the means of iron, manganese, nickel, and soil pH between plots (*P*<0.05; *t*_fe_=2.86; *t*_mn_=5.78; *t*_ni_=-2.89; *t*_pH_=3.37; *df*=16). Iron and manganese average concentrations were higher in Plot 1 (Fig. 2). Nickel average concentration was higher in Plot 2 (Fig. 2). Average soil pH was higher in Plot 1 and lower in Plot 2 (Table 1).

**Table 1.**
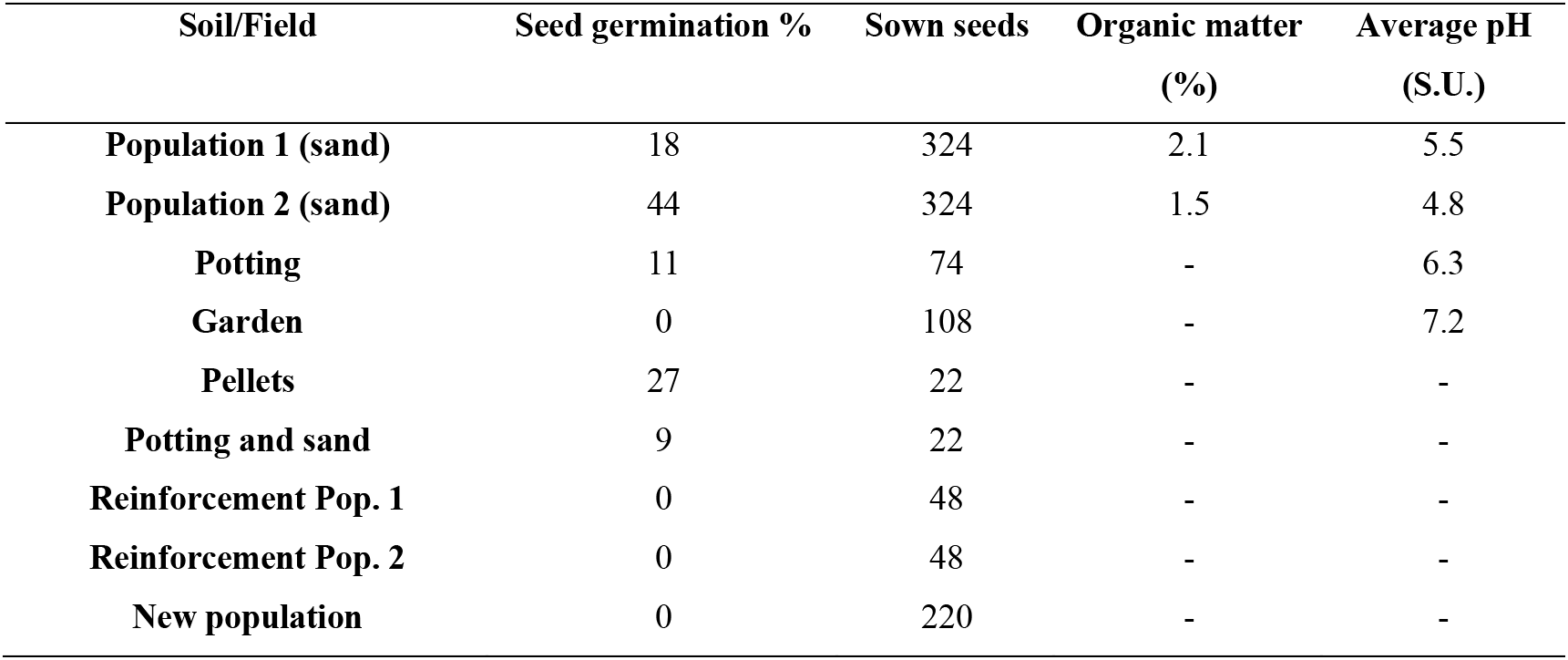
Seed germination percent on different soils and in the field. The total number of seeds sowed, average organic matter, and average soil pH are shown for each case. Dashes were placed when data was not acquired.

**Figure 2.**
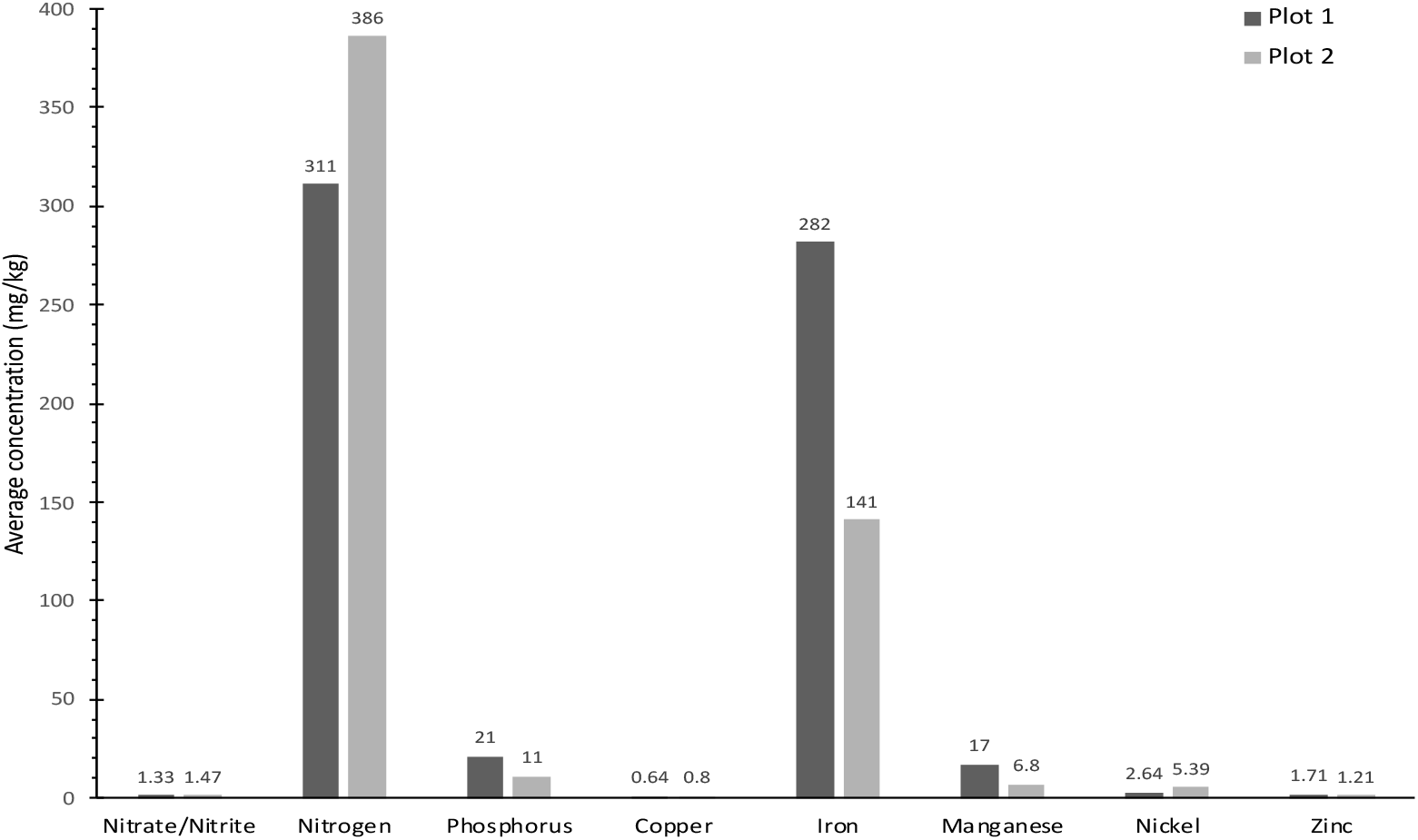
Soil nutrients by plot. Plot 1 was affected by fire. Plot 2 was not affected by fire.

### Plant abundance, reproduction, and seed weight

There was no significant difference in the means of plant abundance between populations in the field (*P*>0.05; *t*=0.4; *df*=3). We found 33 seedlings, 59 juveniles, and 207 adults in Plot 1 for a total of 299 plants. In Plot 2, we found 20 juveniles and 116 adults for 136 plants in total. There was no significant difference in the distribution of the number of seeds per fruit between populations in the field (*P*>0.05; *U*=6041). In our sample, Population 1 produced 393 seeds and Population 2 produced 365 seeds. There was no significant difference in the distribution of seed weight between populations (*P*>0.05; *U*=2513). Average seed weight was 0.0035 g in Population 1 and 0.0034 g in Population 2. We found a significant positive linear relationship between seed weight and average plant height (*P*<0.05; *P*=0.021; *F*=5.64; *t*=2.4; *Pr>t*=0.021; *m*=1353; *R^2^*=0.067; *df*=64; Fig. 3). As a result, plant height increased with seed weight.

**Figure 3.**
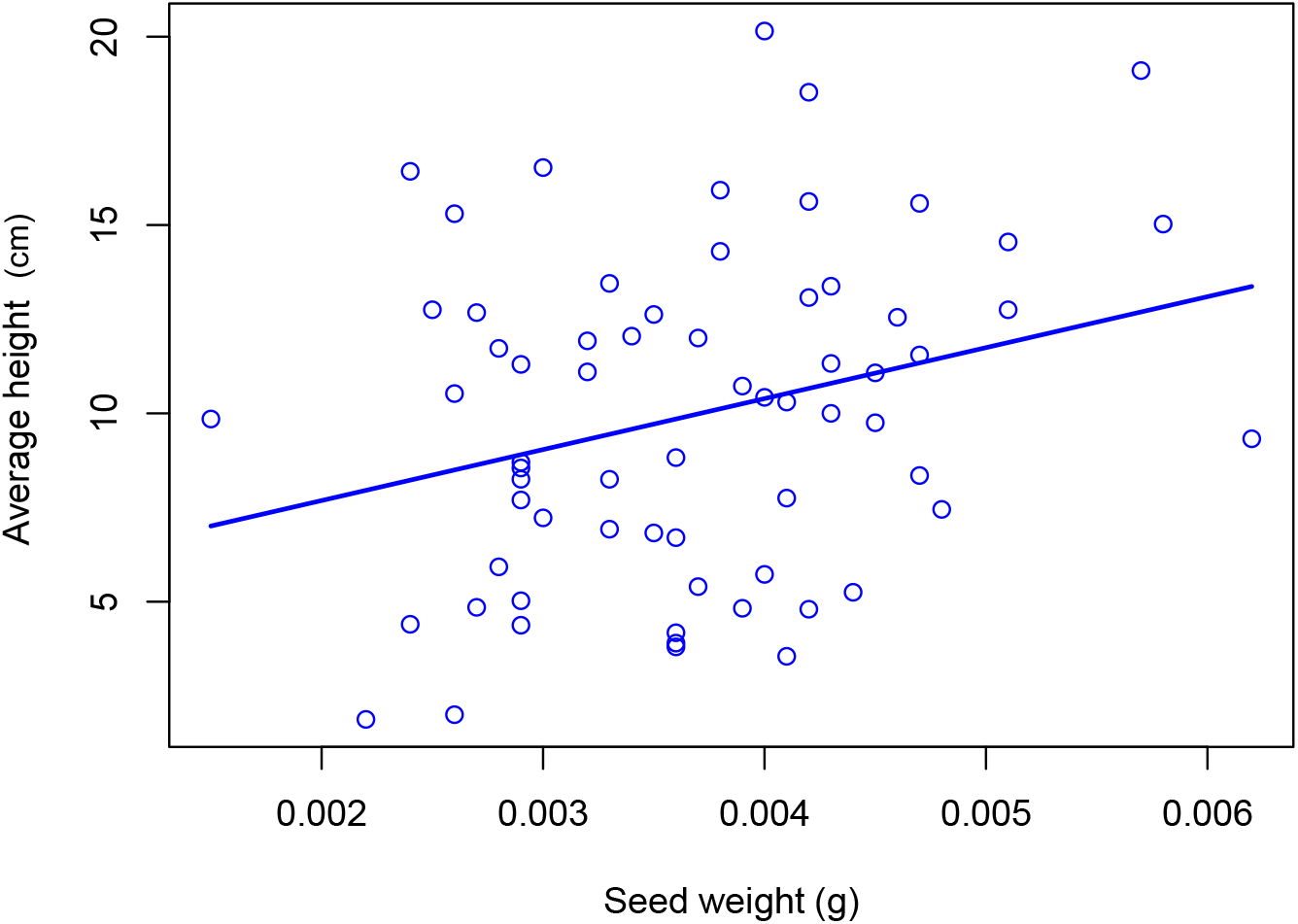
Positive linear relationship between seed weight and plant height.

### Seed germination

Seed germination was achieved by the extension of the radicle and followed by the emergence of cotyledons and the decay of the seed coat (Fig. 1c). Seeds germinated in the seed starting systems (Fig. 1b). Laboratory conditions used for seed germination and plant growth were found to be effective. Air temperature was registered at a minimum of 28°C and a maximum of 31°C. Relative humidity was registered at a minimum of 49% and a maximum of 52%. Soil temperature ranged between 31°C and 34°C. Finally, the water pH was measured at 7.63.

Heavy seeds germinated and lightweight seeds did not. Seeds that germinated weighted between 0.0021-0.0051 g, and seeds that did not germinated weighted between 0.0004-0.0020 g. There was no significant difference in the distribution of germinated seeds by days among populations in the laboratory (*P*>0.05; *U*=1497). Seed germination began in day 2, and the highest germination was achieved in the first 8 days (Fig. 5). In general, most seeds germinated in the first 3 weeks. The seed germination percent was different between populations (Table 1). Population 2 with 44% had a higher germination capacity.

**Figure 5.**
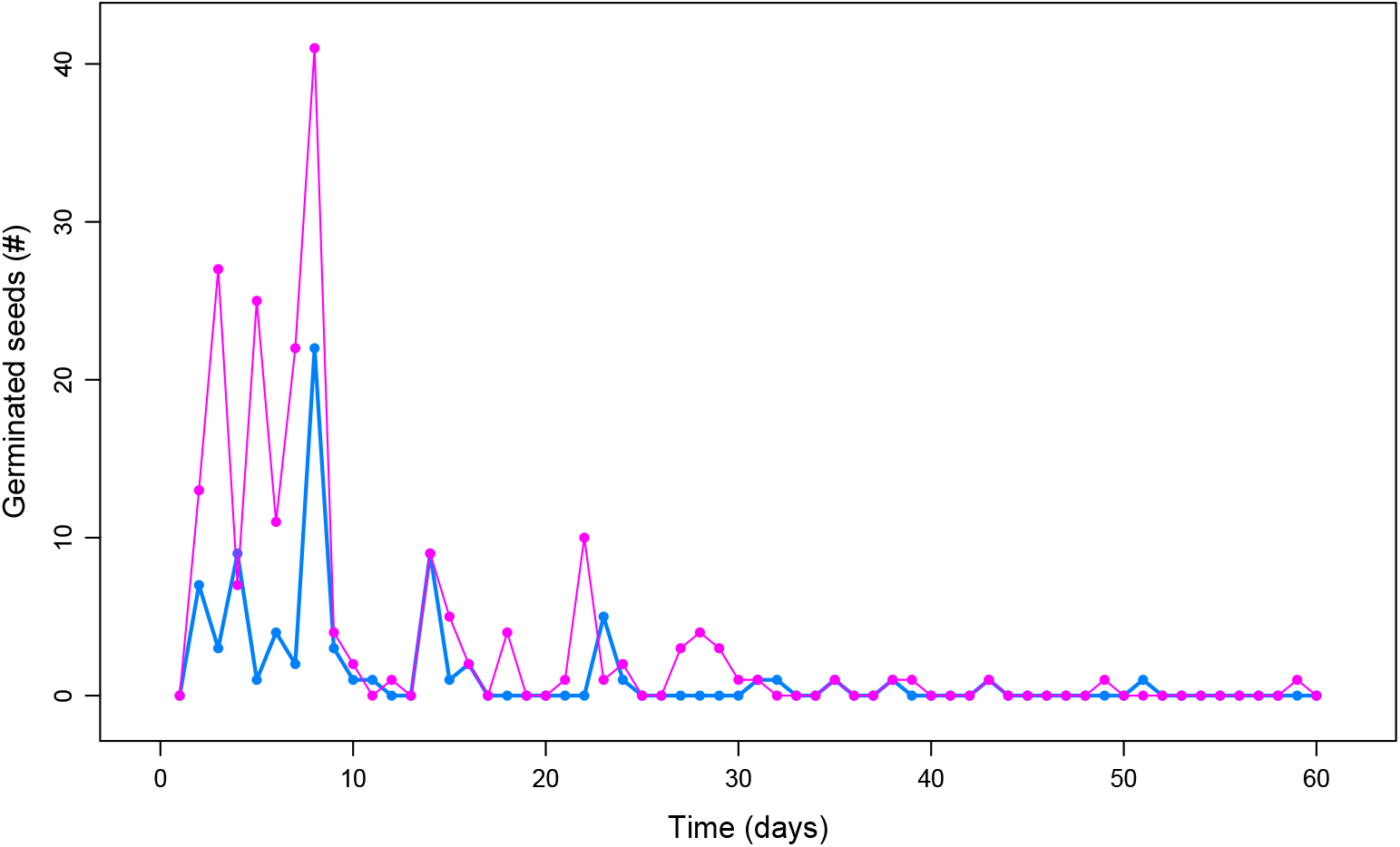
Seed germination rate by population. Population 1, the experimental group, is shown in blue. Population 2, the control group, is shown in pink.

The seed germination percent in different soils was variable. Seeds germinated in potting soil, pellets, and potting mix and sand, but did not germinated in garden soil (Table 1). Field germination percent near Plot 1 at November resulted in 0% and near Plot 2 and in the new area in March resulted in 0% (Table 1). After 5 months, no seed germinated in the field. Instead, other species germinated and grew in the plots. We found a significant negative linear relationship between average soil pH and seed germination percent (*P*<0.1; *P*=0.055; *F*=17; *t*=−4.1; *Pr>t*=0.055; *m*=-17; *R^2^*=0.84; *df*=2; Fig. 6). The seed germination percent increased when average soil pH decreased.

**Figure 6.**
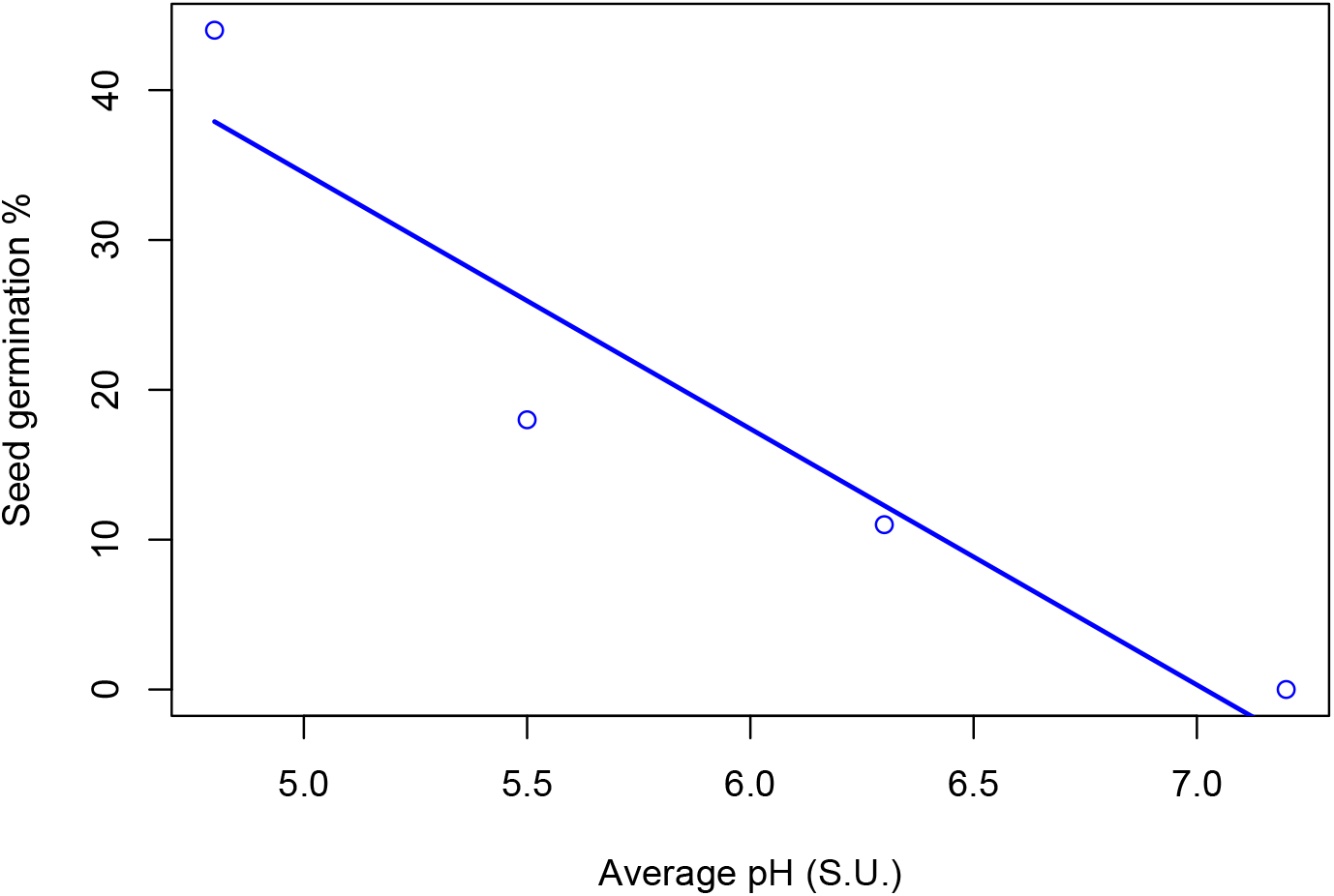
Negative linear relationship between soil pH and seed germination percent.

### Plant growth

Plants grew in the seed starting systems (Fig. 1b). There was a significant difference in the means of plant height between the experimental and control groups in the laboratory (*P*<0.05; *t*=2.74, *df*=190). Plant height increased with time in average and was higher in Population 1, the experimental group (Fig. 7). In general, plants doubled the average height of half time in week 8.

**Figure 7.**
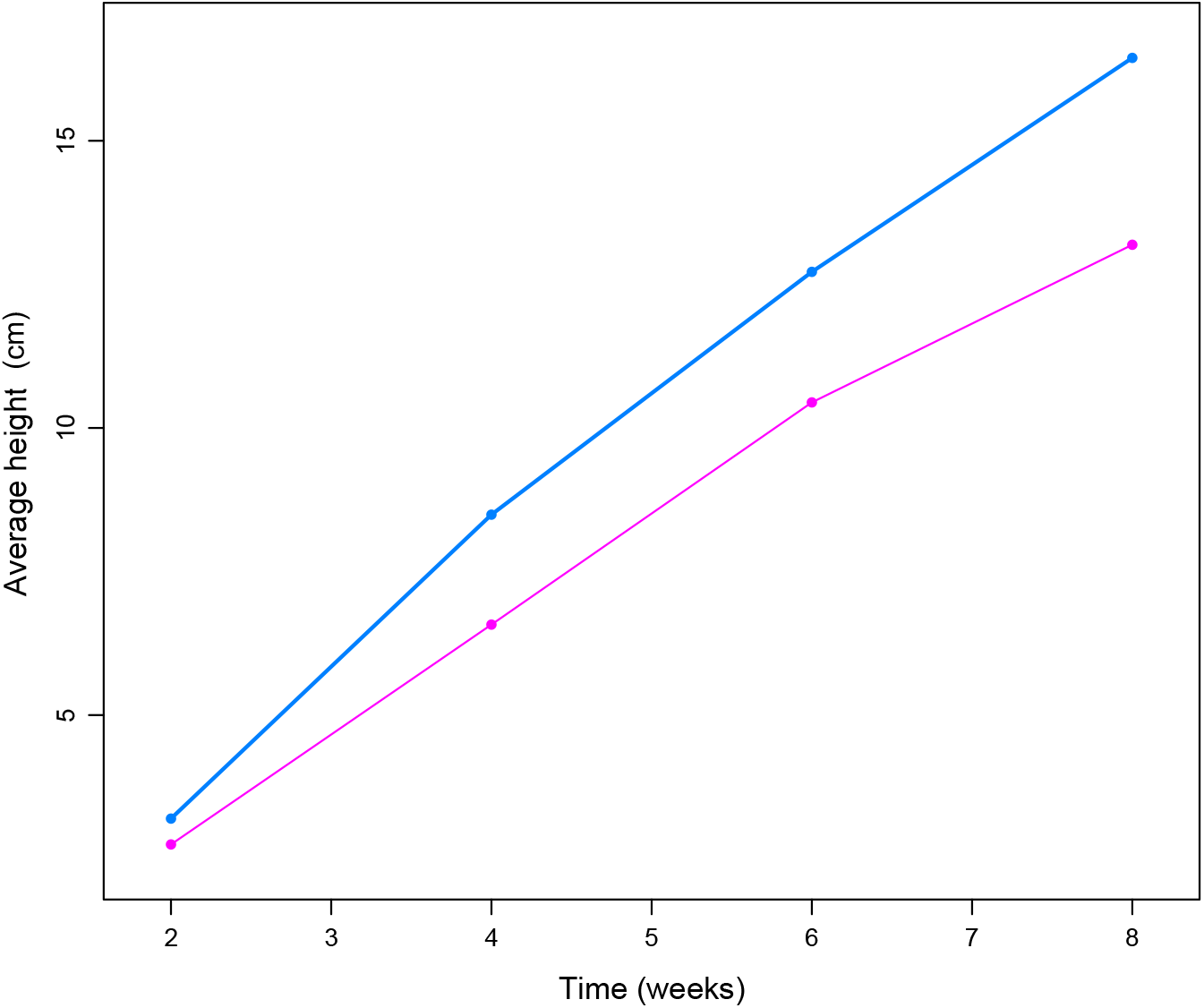
Plant growth by population. Population 1, the experimental group, is shown in blue. Population 2, the control group, is shown in pink.

### Acclimation and reintroductions

Plants acclimated to the semi-natural conditions in the nursery environment. We observed plant mortality in the laboratory before acclimation and in the nursery during acclimation. Field transplantation was possible from plants grown in the laboratory and nursery. Plants reintroduced in populations for reinforcement had variation in survival rate. The survival percent of reintroduced plants in Population 1 started high in the first months, but survival decreased to 0% in the last 2 months. In contrast, reintroduced plant survival (33%; 1 plant) was constant in Population 2 during the 5 months. Reintroduced plants acclimated to the natural conditions and achieved reproductive maturity through time. The percent of plants reproducing in the first two weeks reached 80% and then decreased. Finally, the percent of plants producing buds, flowers, and fruits increased to 80% at week 10 and remained constant until week 12.

## Discussion

### Soil chemistry

The fire caused a significant increase in soil pH, iron, and manganese and a decrease in nickel after 2 years of the fire. Our hypothesis was supported by the increase in soil pH, iron, and manganese in Plot 1. The other nutrients remained similar between plots suggesting leaching in Plot 1. Nutrients like nitrogen and phosphorus can be leached by rainfall and lost from the burned ecosystem decreasing to pre-fire levels (Kutiel and Naveh 1987). Fires have been found to significantly increase soil pH (Kutiel and Naveh 1987, Norouzi and Ramezanpour 2013, Amoako and Gambiza 2019), specifically, high intensity fires significantly increase soil pH (Bridges et al. 2019).

Higher concentrations of iron and manganese and a lower level of nickel in the burned environment could be due to the increased soil pH value. Changes in soil micronutrients can be caused by high intensity fires and can be related to soil pH (Norouzi and Ramezanpour 2013). In agreement, reducing soil pH increases nickel accumulation (Nkrumah et al. 2019). Results have further suggested that burning enhances the availability of some soil nutrients on topsoil for a short period being an unsustainable way of increasing soil productivity (Amoako and Gambiza 2019). Besides fire, agricultural lime can be used to increase soil pH and soil nutrient availability (Ritchey et al. 2016).

### Plant reproduction and seed weight

Results showed no effect of fire in reproduction, seed weight, seed germination, and plant abundance. Contrary to our results, seeds number and seed weight of a wheat species were higher in the recently burned soil (Kutiel and Naveh 1987). For only some species, fruit set was significantly greater in the burned environment (Paritsis et al. 2006). Moreover, fire intensity was found to increase the fruit production of *Chamaecrista keyensis* at short term (Liu et al. 2005). The no effect of fire in the study species is understandable because legumes have shown to be resistant to fire (Fidelis et al. 2016). These show that fire effects on plants, especially reproduction, can be variable depending on the species, fire intensity and frequency, nutrient availability, and aboveground competition (Liu et al. 2005).

### Seed germination

Germination and propagation from seeds were possible in laboratory conditions.

*Chamaecrista glandulosa* var. *mirabilis* may require temperatures around 31-34 °C and high soil moisture for seed germination. The germination percent in the field was 0% in November and March. This suggests that natural conditions were not adequate for germination during these months and germination possibly occurs in the field close to August or September where conditions are favorable. Distinct effects of heat on germination rates have been found in Fabaceae species (Silveira and Overbeck 2013). Environmental factors related to the breaking of dormancy (e.g., temperature fluctuations, soil moisture) might play a more important role in seed germination than the insufficient direct effect of fire-related high temperatures exposure (Fidelis et al. 2016).

Seed germination was related to soil pH in the species, as we previously hypothesized. Soil pH explained 84% of the variance of seed germination percent. This result indicates that seed germination is highly predicted by soil pH, suggesting that *C. glandulosa* var. *mirabilis* is adapted to acid silica sand soils. Germination responds with a high percent at the acid low pH of silica sand soil and germination does not occur at the neutral pH of garden soil. This finding is consistent with previous results where grass species seeds showed better germination towards the acidic range (Singh et al. 1975). This may be because the acidic medium favors the synthesis, action (Singh et al. 1975), and/or function of enzymes (Gafar et al. 2018) necessary for germination. Although there was no significant difference in the seed germination rate, our results of seed germination predictions by soil pH and a higher germination capacity at 4.8 pH suggest that large increases towards neutral pH caused by fires could reduce or inhibit seed germination in the species.

### Plant growth and size

Results showed an effect of post-fire soil nutrients on the species, partially supporting our hypothesis. Greater plant height through time in the experimental group, Population 1, could have resulted because of higher iron and manganese and lower nickel concentrations in the soil. Iron deficiency can affect plants (Norouzi and Ramezanpour 2013), so iron and manganese sufficiency can positively affect plant growth. Also, reducing soil pH increases nickel in plants and decreases shoot biomass and growth (Nkrumah et al. 2019). Studies have reported that natural charcoal and a short-term flush of mineral elements in the soil caused by fire is followed by a positive effect on the growth of herbaceous plants (Kutiel and Naveh 1987, Gundale and DeLuca 2006). Our results confirm that fires significantly increase soil pH and cause changes in soil micronutrients that can significantly increase plant growth.

Plant height was found to be positively related to seed weight, consistent with our hypothesis. This result is in line with previous studies that have found a positive correlation between seed weight and plant size (Hendrix 1984) and a positive correlation between seed mass and shoot length (Mukherjee et al. 2019). Heavy seeds having big embryos and endosperms can produce higher plants. Large seeds may have larger embryos resulting in bigger seedlings (Xu et al. 2016). As a management strategy, heavy seeds can be used for propagation having a higher germination percent (Anjusha et al. 2015, Xu et al. 2016) and producing bigger plants that can have high survival (Schreeg et al. 2005, Dollard 2018) and reproduction (Rojas-Sandoval and Meléndez-Ackerman 2011).

### Reintroductions

Species reintroductions could be done in both populations. Reintroduced plants were able to survive and reproduce through time in the natural habitat. Their survival decreased to 0% in Population 1 due to natural disturbances. Mortality in the first reintroduction in Population 1 was caused by Hurricane María in September 2017. Deaths were due to an uprooted large canopy tree that crushed and bended the plants, as happened for another shrub species in response to a hurricane (Pascarella 1998). Mortality in the second reintroduction in Population 1 was caused by a horse grazing and trampling in the area. This same threat was detected in other study along sheep, cattle, and pig grazing (Fenu et al. 2011).

### Future research and management

Future research could consider experimental species reintroductions. Conservation genetics can be incorporated to develop strategies for the preservation of genetic diversity and ultimately the conservation of biodiversity (Pertoldi et al. 2007). Studies could be designed to test different temperature, humidity, and pH treatments for optimal seed germination and plant growth in the laboratory and field. The development of fire control lines and horse exclusion fences in the natural reserve, landscape maintenance, invasive species prevention, control and eradication, and monitoring are management strategies recommended. Finally, controlled fires can be utilized in vegetated areas for species reintroductions to achieve the preservation of the species.

## Conclusions

In conclusion, large size plants are produced from high weight seeds. *Ex situ* propagation from seeds was an efficient method for the *ex situ* conservation of *C. glandulosa* var. *mirabilis*. Augmentation of populations was a viable technique following *ex situ* conservation. *Ex situ* conservation and species reintroduction were feasible conservation strategies for the species that should be integrated with *in situ* conservation. Species recovery actions, ecological restoration (Heywood 2017), threats control, and landscape conservation are *in situ* conservation strategies that can be implemented to protect and preserve the species and its ecosystem.

## Acknowledgements

We thank Ana G. Méndez University for its laboratory and materials. Specially, we are grateful with Julita Rivera, Dawin Santiago, and Armando Canchani for their support in the laboratory. Also, we are grateful with Christian Velez, MSEM and José J. Fumero, Ph.D. for the manuscript revision. Finally, we acknowledge the Department of Natural and Environmental Resources of Puerto Rico for access to RNLT and the species (permit number: 2014-EPE-031).

## Authors Contribution

Conceptualization, F.J.V.T. and J.A.L.C.; Methodology, F.J.V.T. and J.A.L.C.; Investigation, F.J.V.T. and J.A.L.C.; Data Curation, F.J.V.T.; Software, F.J.V.T.; Validation, F.J.V.T.; Formal Analysis, F.J.V.T.; Visualization, F.J.V.T.; Writing – Original Draft, F.J.V.T.; Writing – Review & Editing, F.J.V.T. and J.A.L.C.; Resources, J.A.L.C.; Supervision, J.A.L.C.; Project Administration, J.A.L.C.

